# Postbiotic Binding of Micro- and Nanoplastics: In Vitro Intestinal Epithelial Protection and Proof of Concept in the Human Mouth

**DOI:** 10.64898/2026.05.11.724280

**Authors:** Eva A. Berkes, Oded Oron, Adriana K. Wood, Phineas N. Monsul, Nicholas T. Monsul

## Abstract

Micro- and nanoplastics (MNPs) are now recognized as ubiquitous dietary and environmental contaminants, yet practical strategies to reduce gastrointestinal exposure remain limited. This study evaluated whether Qi601, a heat-inactivated *Limosilactobacillus fermentum* biofilm-derived postbiotic, could bind plastic particles and reduce intestinal epithelial plastic burden. Prior probiotic studies have demonstrated live bacterial adsorption of MNPs and mitigation of MNP-associated toxicity in vivo; here, we evaluate whether a nonviable postbiotic preparation can produce analogous MNP-binding and epithelial-protective effects. Qi601 durably bound polystyrene nanoplastics under in vitro simulated digestion conditions. In Caco-2 intestinal epithelial monolayers, Qi601 reduced surface-associated and intracellular nanoplastic burden in both protection and rescue models, indicating decreased epithelial particle interaction both before and after established nanoplastic exposure. Multimodal imaging, including confocal microscopy, atomic force microscopy, and scanning electron microscopy, confirmed close physical association between Qi601 and nanoplastics. Finally, a first-in-human proof-of-concept chewing-gum study showed Qi601 binding in the human mouth to heterogeneous gum-derived microplastic fragments released during mastication. Together, these findings support the concept of postbiotic intervention for gastrointestinal epithelial protection against ingested MNPs.

## Introduction

Over the past century, global plastic production has grown rapidly from essentially zero to hundreds of millions of tons per year. More than half of all plastic ever manufactured has been produced since 2000, meaning that baseline exposure levels for the current population is substantially higher than for any prior generation.^1^ The durability of plastic results in the long-term accumulation of plastic waste in the environment, where they fragment into micro- and nanoplastics, collectively referred to here as micronanoplastics, or MNPs.^2,3^

Due to their ubiquitous presence in water, air and food, the accumulation of MNPs has raised growing concern about their potential ecological and human health impacts. Consistent with systemic exposure, MNPs have been detected in many human tissues, including blood, intestinal tract, urine and brain^4–7^. Nanoplastics are of particular concern because their small size may render them able to cross biological membranes^8–10^.

Recommended MNP mitigation measures are primarily environmental, focusing on household and lifestyle modifications. Nascent research regarding biological mitigation suggests that dietary fibers such as chitosan and wheat bran, plasma apheresis and microbiome-mediated plastic degradation may have potential to decrease human MNP burden^11–14^. Recent studies of probiotics and their metabolites also show promise as dietary interventions by physically binding plastics to decrease their uptake (probiotics), or to ameliorate MNP-associated toxicity (probiotics and metabolites) ^15–20^. For example, live LAB strains have been shown to adsorb micro- and nanoplastics, promote fecal elimination, reduce tissue retention, and improve gut-barrier or inflammatory endpoints in murine models. However, live-probiotic approaches have several limitations when considered as MNP mitigants: variable gastrointestinal survival, host-to-host variability, uncertain persistence, shelf-stability constraints, MNP toxicity to organisms such as lactic acid bacteria (LAB) and the theoretical possibility that live organisms could retain bound particles near the mucosal interface rather than favor luminal clearance. ^21–24^

Postbiotics, defined as preparations of inanimate microorganisms and/or their components that confer a health benefit, may offer advantages in this context because they do not require microbial viability^21^. Although live probiotic strategies for MNP adsorption and toxicity mitigation are emerging, little is known about whether postbiotic microbial material can function as a luminal binding platform for MNPs under human-relevant digestive conditions. This study aims to help address that gap by evaluating the influence of a postbiotic LAB preparation, “Qi601”, on plastic particle behavior in models relevant to the human gastrointestinal environment.

Because MNPs undergo physicochemical alterations during gastrointestinal transit which can influence their biological activity and toxicity^22^, an adapted simulated human digestion model (International Network of Excellence on the Fate of Food in the Gastrointestinal Tract, INFOGEST) was used to produce digestates containing NPs with Qi601 postbiotic, and digestates evaluated in a polarized model of the human intestinal barrier^23,24^. Cellular adhesion and intracellular uptake were localized and quantified using confocal laser scanning microscopy (CLSM), and scanning electron microscopy (SEM) and atomic force microscopy (AFM) used to visualize microstructural interactions between NP and Qi601 postbiotic. Finally, a first-in-human proof-of-concept study evaluated oral binding of Qi601 to microplastics using chewing gum as the microplastic-releasing vehicle. We hypothesized that Qi601 would physically bind MNPs during simulated digestion, retain bound particles within a luminal/postbiotic phase, and thereby reduce epithelial surface association and intracellular particle burden. To test this, we combined adsorption assays, simulated digestion, Caco-2 protection and rescue models, transfer-stability testing, multimodal microscopy, and a human oral chewing-gum proof-of-concept study.

## Materials and Methods

### Postbiotic strain and postbiotic preparation

The postbiotic Qi601 material was provided by Quorum Innovations Inc (Sarasota, FL). The strain in this study, *Limosilactobacillus fermentum LfQi6*, has been determined as Generally Recognized as Safe (GRAS) for its intended use. The strain was grown as a biofilm culture, centrifuged, washed, heat-inactivated and lyophilized using proprietary methodology in a cGMP-registered facility^25^. The resulting food-grade postbiotic biofilm cellular mass (“Qi601”) lyophilate was used for subsequent experiments.

### In vitro optimization of postbiotic dose for maximum PSNP adsorption

The Qi601 dose associated with maximal adsorption of 100-nm fluorescent polystyrene nanoplastics (PSNP) was determined using an in vitro adsorption assay. Qi601 doses ranging from 0.5–10 mg were suspended in 900 µL PBS, followed by addition of 10 µL PSNP diluted 1:1 in PBS from the manufacturer-supplied concentrate (Fluoresbrite™ YG Microspheres, 0.1 µm; Polysciences, Warrington, PA; #07150). Each condition was performed in triplicate, with PSNP-only, Qi601-only, and PBS-only controls.

Samples were incubated for 1 h at 37°C with shaking at 250 rpm, then centrifuged at 1,700 × g for 10 min. Supernatants were transferred to black 96-well plates and fluorescence was measured using a Synergy LX multimode microplate reader with a 485/528 nm filter set and gain of 50. Background-corrected supernatant fluorescence was used to quantify unbound PSNP, with lower supernatant fluorescence indicating greater PSNP adsorption to Qi601. Mean fluorescence, standard deviation, and percent adsorption were calculated for each condition relative to PSNP-only controls. Qi601–PSNP adsorption was calculated as:

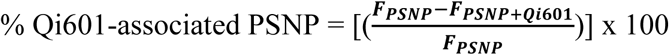

where 𝐹_𝑃𝑆𝑁𝑃_is the background-corrected fluorescence of the PSNP-only supernatant and 𝐹_𝑃𝑆𝑁𝑃+𝑄𝑖601_is the background-corrected fluorescence of the PSNP-plus-Qi601 supernatant after incubation and centrifugation under identical conditions. Based on observed adsorption efficiencies under these conditions, subsequent experiments were performed with 3 mg of Qi601.

### Using simulated digestion to evaluate nanoplastic interactions with Qi601

The static INFOGEST model was adapted to evaluate Qi601–PSNP interactions under simulated gastrointestinal digestion conditions.

One of the most established in vitro digestion methods is the static INFOGEST model, which uses defined digestive-phase conditions to simulate human oral, gastric, and intestinal digestion.²⁶ The INFOGEST model was adapted to evaluate Qi601–nanoplastic interactions under simulated digestion conditions. One hundred-nanometer fluorescent PSNP were incubated with or without Qi601 through sequential oral, gastric, and intestinal phases using commercial synthetic digestive fluids. Volumes were adjusted to maintain fluorescence within assay detection limits, and all conditions were performed in triplicate.

Qi601 was prepared as described above and added to each tube at 100 µL per tube. For the oral phase, samples were incubated with 200 µL artificial saliva (AS; Pickering Laboratories, CA, USA; pH 6.8; #1700-0304) and 10 µL of 100-nm PSNP diluted 1:1 v/v in PBS for 3 min at room temperature. Fed-state simulated gastric fluid was then added (800 µL; FeSSGF; BioChemazone, Alberta, Canada; #BZ415), and samples were incubated at 37°C for 1 h with shaking at 250 rpm. Artificial intestinal fluid was subsequently added (800 µL; AIF; BioChemazone; #BZ176), followed by a second 1-h incubation at 37°C with shaking at 250 rpm. Parallel assays were performed in each simulated digestive fluid alone to assess effects of each digestive fluid on Qi601–PSNP interactions.

Following digestion, samples were centrifuged as described above to pellet Qi601-associated material. Supernatants were collected without disturbing the pellet, transferred to fresh tubes, and plated in black 96-well plates in technical triplicate. Fluorescence was measured as described above, and background-corrected fluorescence was used to calculate PSNP removal from the supernatant relative to PSNP-only controls. Digestates were frozen at −20°C for subsequent Caco-2 experiments.

### Generation of in vitro human intestinal model

Caco-2 cells (HTB-37™; American Type Culture Collection, VA, USA) were used for in vitro PSNP uptake experiments. Caco-2 is a human colorectal adenocarcinoma-derived epithelial cell line commonly used as an intestinal epithelial model. Cells were maintained in T-75 culture flasks (Corning, #430641) in complete Eagle’s Minimum Essential Medium (EMEM; American Type Culture Collection, VA, USA; #30-2003) supplemented with 20% v/v fetal bovine serum (FBS; Gibco, Thermo Fisher Scientific, MA, USA; #A5256701) and 1% penicillin-streptomycin solution, 10,000 U/mL (Gibco, Thermo Fisher Scientific; #15140122).

Cells were incubated at 37°C in a humidified atmosphere containing 5% CO₂. Medium was refreshed every 2 days until cells reached approximately 50% confluence, after which medium was refreshed daily until cells reached 80–95% confluence. For subculturing and experimental seeding, cells were trypsinized at 37°C with 0.25% trypsin/2.21 mM EDTA (Corning, NY, USA; #25-053-CI) and neutralized with an equal volume of trypsin neutralizer (Gibco, Thermo Fisher Scientific; #R002100). The cell suspension was centrifuged at 400 × g for 10 min, and the cell pellet was resuspended in 1–2 mL complete medium.

For experiments, cells were seeded into flat-bottom 24- and 96-well plates (Costar, Corning, NY, USA; #3524 and #3595) at 2 × 10⁴ cells/well. Before counting and seeding, cell suspensions were mechanically triturated by passage through a 25-gauge needle (BD PrecisionGlide™ 1 inch; Becton Dickinson; #305125) to minimize cell aggregation and spheroid formation. Experiments were performed once cells reached approximately 85% confluence.

### Assessment of Caco-2 monolayer tolerance to simulated digestive fluids

To align Caco-2 exposure conditions with the adapted INFOGEST model, experiments were conducted using a digestive-fluid mixture (DF) composed of artificial saliva (AS), fed-state simulated gastric fluid (FeSSGF), and artificial intestinal fluid (AIF). Because these fluids are nutrient-deficient and contain digestive enzymes, baseline Caco-2 tolerance to DF exposure was first evaluated. Cells were seeded in complete EMEM at 2 × 10⁴ cells/cm² in 24-well plates and maintained at 37°C in a humidified 5% CO₂ atmosphere until approximately 85% confluent, with medium changes as described above.

Before cell exposure, DFs were heat-inactivated at 90°C for 5 min to reduce enzymatic activity and then filtered through 0.22-µm PES syringe filters (Genesee Scientific, CA, USA; #60-1850). Cells were exposed to undiluted DF or DF diluted to 75%, 50%, or 25% v/v in complete EMEM. These diluted digestive fluid/media mixtures are referred to as DF/EM. Control wells were maintained in complete EMEM alone. To identify the highest DF concentration compatible with Caco-2 monolayer integrity, cell adherence and morphology were assessed by light microscopy at 2, 5, 9, and 24 h after exposure.

### Rescue assay: Qi601-mediated reduction of Caco-2 surface-associated and intracellular PSNP

The Caco-2 model was used to evaluate whether Qi601 could reduce PSNP burden after PSNP had already associated with and entered intestinal epithelial cells (“rescue assay”). Caco-2 cells were cultured in 24-well plates in complete EMEM until approximately 85% confluent. On the day of the experiment, cells were washed twice with PBS and exposed to 0.1% PSNP diluted in DF/EM, with 1 mL applied per well. Cells were incubated for 24 h at 37°C in a humidified 5% CO₂ atmosphere. The following day, PSNP-containing medium was aspirated, and cells were washed twice with PBS to remove unbound PSNP. Qi601 postbiotic was then applied at 3 mg/mL in DF/EM, and cells were incubated for an additional 5 or 24 h. Controls included untreated cells, Qi601-only cells, and PSNP-only cells.

At each time point, surface-associated and intracellular PSNP fractions were measured using a protocol adapted from Dos Santos et al., who optimized washing conditions to remove surface-bound PSNP before quantifying remaining intracellular/cell-associated signal. Briefly, Caco-2 cells were washed three times with PBS to remove Qi601, loosely associated PSNP, and spent DF/EM. To recover the extracellularly recoverable surface-associated fraction, monolayers were incubated with 600 µL PBS containing Ca²⁺ and Mg²⁺ for 4 min at room temperature. The resulting surface-wash supernatant was collected for fluorescence analysis. Cells were then lysed with pre-warmed 0.5% Triton X-100 in PBS, centrifuged as before and lysates supernatants collected for quantification of intracellular/cell-associated PSNP signal and protein quantification.

Protein content per well was measured using the Pierce™ BCA Protein Assay Kit (Thermo Fisher Scientific; #23225). For each sample, 200-µL technical replicates were transferred to black flat-bottom microtiter plates, and fluorescence was measured as described above. Fluorescence values were background-corrected and normalized to total protein per well.

Surface-associated and intracellular PSNP signal per milligram of cellular protein were calculated as:

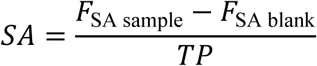

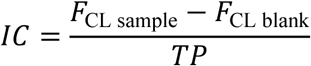

where 𝐹_SA sample_is fluorescence recovered from the surface-associated fraction, 𝐹_SA blank_is fluorescence recovered from the corresponding negative-control surface wash, 𝐹_CL sample_is fluorescence recovered from the cell lysate fraction, 𝐹_CL blank_is fluorescence recovered from the corresponding negative-control lysate, and 𝑇𝑃is total cellular protein per corresponding well.

### Protection assay: Qi601-mediated protection against Caco-2 PSNP adhesion and uptake

Caco-2 cells were cultured in 24-well plates until approximately 85% confluent. On the day of experiment, the cells were washed twice with PBS and treated with Qi601, 3 mg/ml, in 500 µl of DF/EM. Background control wells received DF/EM only. Qi601 was incubated for 30 minutes at 37°C in a humidified atmosphere with 5% CO₂. Following pre-incubation, 0.1% PSNP were added to the appropriate wells in 500 µL DF/EM. Cells were then incubated for 5 or 24 hours at 37°C in a humidified atmosphere containing 5% CO₂. Surface-associated and intracellular PSNP fractions were collected and quantified as described above.

### Epithelial transference: Qi601-associated PSNP transfer to Caco-2 cells

Caco-2 monolayers were used to determine whether PSNP bound within Qi601–PSNP agglomerates (QPAs) could become available for epithelial surface association or intracellular uptake after simulated digestion. Caco-2 cells monolayers were cultured in 24-well plates until approximately 85% confluent. On the day of experiment, postbiotic Qi601 was suspended at 3 mg/ml in DF/EM and co-incubated with selected PSNP concentrations, 0.1%, 0.5%, 1% and 2.5%, for 1 hour at 37°C with shaking at 250 rpm. Following incubation, samples were centrifuged at 1,700 x g for 10 minutes. The postbiotic-containing pellet was retained. The unbound PSNP in the supernatant were measured with the Synergy LX multimode microplate reader to validate the expected adsorption by the QPAs, and downstream transference calculation. The pellet was washed three times in PBS, resuspended in DF/EM to a final Qi601 concentration of 3 mg/ml, and applied to Caco-2 monolayers for 5 and 24 hours at 37°C in a humidified atmosphere containing 5% CO₂.

PSNP-only comparator controls were prepared based on the quantity of PSNP fluorescence associated with Qi601 at each input PSNP concentration. Qi601-associated PSNP fluorescence was calculated by subtracting the fluorescence remaining in the post-centrifugation supernatant from the total input PSNP fluorescence. PSNP-only comparator wells were then prepared using free PSNP at fluorescence levels equivalent to the calculated Qi601-associated PSNP fluorescence for each condition.

Apparent PSNP transfer from QPAs to Caco-2 cells was calculated at each PSNP concentration using the following equation, and results were reported as percentages relative to PSNP-only controls.

The percentage of Qi601-associated PSNP fluorescence transferred to Caco-2 cells was calculated as:

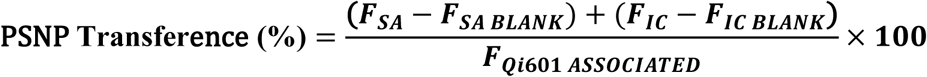

Where:

𝐹_𝑆𝐴_ = fluorescence recovered from the surface-associated fraction

𝐹_𝑆𝐴 𝐵𝐿𝐴𝑁𝐾_ = fluorescence recovered from the corresponding membrane-bound negative control

𝐹_𝐼𝐶_ = fluorescence recovered from the intracellular fraction

𝐹_𝐼𝐶 𝐵𝐿𝐴𝑁𝐾_ = fluorescence recovered from the corresponding intracellular blank control fraction

𝐹_𝑄𝑖601 𝐴𝑆𝑆𝑂𝐶𝐼𝐴𝑇𝐸𝐷_ = fluorescence associated with Qi601 pellet after loading and washing, before Caco-2 exposure

After 5- or 24-hours exposure, membrane-associated and intracellular PSNP fluorescence were quantified as described above.

### Confocal laser scanning microscopy

Caco-2 cells were cultured as monolayers in 8-chamber MilliporeSigma™ Millicell™ EZ Slide microscope slides (Fisher Scientific, MA, USA; #PEZGS0816) in complete EMEM until approximately 85% confluent. Cells were then exposed to 300 µL of 0.1% PSNP diluted in DF/EM and incubated for 24 h at 37°C in a humidified atmosphere containing 5% CO₂. The following day, chambers were washed to remove spent medium and unbound PSNP. Selected chambers were then treated with 300 µL Qi601 postbiotic at 3 mg/mL in DF/EM for either 5 or 24 h. Controls included PSNP-only exposed cells, Qi601-only exposed cells, and untreated cells to account for background fluorescence.

At the end of each time point, cells were rinsed three times with PBS to remove Qi601 and spent medium. In one chamber that had received both PSNP and Qi601, only one PBS wash was performed to intentionally retain a portion of Qi601 material for qualitative assessment of PSNP adsorption by Qi601 on the Caco-2 monolayer.

Cells were fixed with 4% paraformaldehyde (Thermo Scientific; #AAJ60401AP) for 20 min at room temperature, washed three times with PBS to remove excess fixative, and permeabilized with 0.1% Triton X-100 in PBS. F-actin was stained with rhodamine-phalloidin diluted 1:40 in PBS for 20 min at room temperature. Nuclei were stained with 200 µL DAPI at 0.1 µg/mL (Biotium, CA, USA; #40043) diluted in PBS for 10 min at room temperature. To visualize Qi601 material remaining on the Caco-2 monolayer, extracellular carbohydrate-rich matrix components were stained with fluorescently labeled concanavalin A (ConA-CF™ 350 conjugate; Biotium, CA, USA; #29137) diluted to 200 µg/mL in deionized water and incubated at 37°C for 15 min. A deionized-water wash was performed between staining steps, and chambers were allowed to air-dry. ProLong™ Gold Antifade Mountant (Invitrogen) was applied, coverslips were placed, and the mounting medium was allowed to cure overnight before imaging.

Fluorescent PSNP and cellular stains were visualized using an Olympus FV1200 spectral inverted laser-scanning confocal microscope. Images were acquired using excitation/emission settings of 330–385 nm/420 nm for DAPI and CF™ 350 ConA, 470–495 nm/510–550 nm for fluorescent PSNP, and 530–550 nm/575 nm for rhodamine-phalloidin. These signals were displayed as blue, green, and red channels, respectively.

Images for assessing Caco-2-associated PSNP burden and cellular localization were acquired from 15–20 regions of interest (ROIs) per condition across three chambers per treatment or control group. Z-stack images were acquired using identical imaging parameters across all groups. Image analysis was performed using ImageJ version 1.54p (National Institutes of Health, MD, USA). PSNP fluorescence intensity was calculated on a per-pixel basis within the segmented cell area or defined field of view. Because discrete particle counts did not account for particle size or fluorescence intensity, the primary outcome was total ROI-associated PSNP fluorescent burden. Total fluorescent area was determined from ROI pixel-area measurements quantified in ImageJ.

### Scanning electron microscopy (SEM)

Samples were washed in buffer, collected on hydrophilic Isopore 0.4 µm 13 mm polycarbonate filters (MilliporeSigma, MA, USA; #HTTP01300), fixed in 2.5% glutaraldehyde, dehydrated through a graded ethanol series, critical-point dried, mounted on aluminum stubs, sputter-coated with gold/palladium, and imaged by scanning electron microscopy (SEM).

### Atomic force microscopy (AFM)

Qi601 stock suspension at OD600 4.0 was diluted 1:200 v/v in water and incubated with PSNP diluted 1:1,000 v/v in water for 1 h. The supernatant containing unbound PSNP was removed, and the sediment containing Qi601-associated PSNP was rinsed. Twenty-microliter aliquots of rinsed sediment diluted 1:100,000 v/v in water were dried onto clean, uncoated microscope slides. Qi601-only, PSNP-only, and Qi601-plus-PSNP samples were imaged by atomic force microscopy using an nGauge AFM (ICSPI Corp., ON, Canada).

### Human oral proof-of-concept of Qi601 interaction with microplastics: chewing gum study

To complement the in vitro PSNP-binding and intestinal epithelial cell-association assays, a chewing gum study using human participants was performed to assess whether Qi601–plastic interactions could occur under physiologic salivary and mastication conditions. This experiment was designed as a qualitative imaging-based proof-of-concept study rather than a quantitative exposure assessment. Chewing gum was selected as a reproducible, common oral exposure model because gum-base material can release microplastic particles during mastication. Based on prior studies demonstrating microplastic release from chewing gum in human saliva^26^, this study was designed to evaluate whether Qi601 could associate with gum-derived microplastics generated during chewing. The study was involved adult volunteers in a minimal-risk chewing-gum saliva-collection study under ordinary food-use conditions after providing informed consent. Specimens were deidentified, no personal health information was collected, and no diagnostic, genetic, or clinical testing was performed. Prospective IRB review was not required, per legal review.

Efforts were made to minimize exogenous plastic contamination during sample collection and processing. Glass serological pipettes, glass tubes, 100-mL glass beakers, and aluminum foil sheets were used whenever possible. Glassware and aluminum sheets were rinsed three times with deionized water, submerged in 5% Extran® 300 surfactant for 45 min, and rinsed three additional times with deionized water. Materials were then rinsed with 95% ethanol and air-dried under laminar airflow in a biosafety cabinet. Once dry, glassware was covered with pre-treated aluminum sheets. All liquid reagents used for saliva sample processing were stored in decontaminated glassware. Participants were instructed to avoid eating, drinking, and oral hygiene activities for at least 2 h before the experiment, to wear non-synthetic clothing, and to wash their hands thoroughly before sample collection.

### Saliva collection from the gum-chewing experiment

A commercially available chewing gum brand was arbitrarily selected for the experiment. To reduce baseline oral particulate contamination, participants first rinsed their mouths with 10 mL sterile normal saline for 30 seconds, and the rinsate was collected into a glass beaker. Before gum chewing, participants were instructed not to swallow during the collection period and to expectorate saliva directly into the provided glass beaker whenever they felt the urge to swallow. For the gum-only control condition, participants chewed one piece of gum and expectorated saliva until 10 mL was collected. To recover residual gum-derived particles remaining in the oral cavity, participants then rinsed with an additional 10 mL sterile normal saline, which was added to the saliva sample.

For the gum + Qi601 condition, participants chewed a fresh piece of gum for approximately 5 s before 10 mg Qi601 was introduced. This brief initial chewing period was included to allow mechanical release of gum-associated particles before concomitant exposure to Qi601 during continued mastication. Participants continued chewing while expectorating saliva until 10 mL was collected, followed by an additional 10 mL sterile saline oral rinse that was added to the collected sample. This generated a combined sample containing saliva, gum-derived particles, and Qi601. Samples were stored at 4°C until subsequent processing.

### Saliva processing

Glass serological pipettes, glass tubes, and aluminum foil cleaned as described above were used throughout the sample-processing protocol. Where feasible, samples were handled under laminar flow conditions to minimize airborne particulate contamination. During centrifugation, glass tubes were placed inside 15-mL conical tubes to accommodate the centrifuge rotor. Saliva samples were transferred from glass beakers to 25-mL glass tubes and allowed to sediment at room temperature for 30 min. The supernatant was carefully removed, and the sediment-enriched fraction containing Qi601, gum-derived particles, and residual biological material was retained.

Enzymatic processing conditions were established in preliminary method-optimization experiments to reduce salivary organic material while minimizing loss of postbiotic Qi601 during sample preparation. Four milliliters of warmed 0.125% trypsin were added to each tube containing sediment. The trypsin solution was prepared by diluting 0.25% trypsin/2.21 mM EDTA 1:1 in PBS (Corning, USA; #25-053-CI). Samples were transferred to 5-mL glass tubes, covered with foil, and incubated at 37°C for 45 min to reduce proteinaceous and cellular biological material. The tubes were centrifuged at 1,700 × g for 5 min, supernatant was discarded, pellet was washed once with PBS, centrifuged again at 1,700 × g for 5 min, and the wash supernatant discarded.

DNase I treatment was performed to reduce residual extracellular DNA (RNase-Free DNase Set; Qiagen, CA, USA; #79254). DNase I was reconstituted according to the manufacturer’s instructions, and 160 µL of reconstituted enzyme solution was added to each pellet. Samples were incubated at room temperature for 15 min, centrifuged at 1,700 × g for 5 min, and the supernatant was discarded. Pellets were then washed with 500 µL PBS, centrifuged at 1,700 × g for 5 min, and the wash supernatant discarded.

Nile Red was used to identify fluorescent hydrophobic polymer-like particles. Nile Red (Sigma-Aldrich, St. Louis, MO, USA; #72485) was prepared in DMSO and diluted in PBS to a final concentration of 0.005%. Pellets were resuspended in 200 µL 0.005% Nile Red solution, incubated for 5 min at room temperature, and centrifuged at 1,700 × g for 5 min. The Nile Red solution was discarded. Residual dye was removed with a 70% ethanol wash followed by centrifugation at 1,700 × g for 5 min. Samples were then washed once with 500 µL deionized water, centrifuged at 1,700 × g for 5 min, and excess water was removed. Pellets were resuspended in 1 mL deionized water, and each sample was diluted 1:10 in deionized water. A 25-µL aliquot of each diluted sample was mounted on a clean glass slide with a coverslip and examined using an Olympus BX60 fluorescence microscope. Nile Red-stained particles were visualized using 540–550 nm excitation and 572–620 nm emission settings. Particles were interpreted as gum-derived microplastics when Nile Red fluorescence colocalized with UV excited fluorescence and/or polymer-like morphology in gum-exposed samples, relative to baseline saliva controls.

### Red nanoplastics sedimentation by Qi601

Qi601 (33mg) was resuspended in 1 ml of 1XPBS in a 15 ml conical tube, followed by the addition of 100μl of 0.20μm red dye microspheres (Polybead® Polystyrene Red Dyed Microsphere 0.20 μm, Polysciences, Warrington, PA #15705). The negative control was the red dye microsphere resuspended in 1XPBS alone. The mixture was then incubated for 10 minutes at 37°C and shaking at 250RPM. Then, 10 ml of 1X PBS were added to each tube, and the mixtures were inverted a few times to mix the formed QPA and red dye of the negative control to form a homogenous red solution. The samples were left on the benchtop and recorded until the majority of the QPA settled at the bottom of the tube.

### Statistical analysis

Statistical analyses were performed using Microsoft Excel or GraphPad Prism (GraphPad Software, Boston, MA). Data are presented as mean ± SD. Comparisons between treatment groups and corresponding control groups were performed using unpaired, two-tailed Student’s t tests. Comparisons involving multiple treatment groups were analyzed using one-way ANOVA followed by Dunnett’s multiple-comparisons test versus control. Fluorescent microscopy analysis was done with ImageJ, version 1.54p. Statistical significance was defined as p < 0.05. Significance levels were denoted as follows: ns, p > 0.05; *p < 0.05; **p < 0.01; ***p < 0.001; ****p < 0.0001.

## Results

Initial binding showed that PSNP adsorption increased with Qi601 dose, reaching a maximum of 87.6% at 3 mg (Fig. 1A). Adsorption appeared dose-dependent up to 3 mg and then plateaued. This plateau may reflect saturation of detectable particle-binding capacity, incomplete pelleting of Qi601–PSNP aggregates, or reduced assay resolution at higher postbiotic masses.

**Figure 1.**
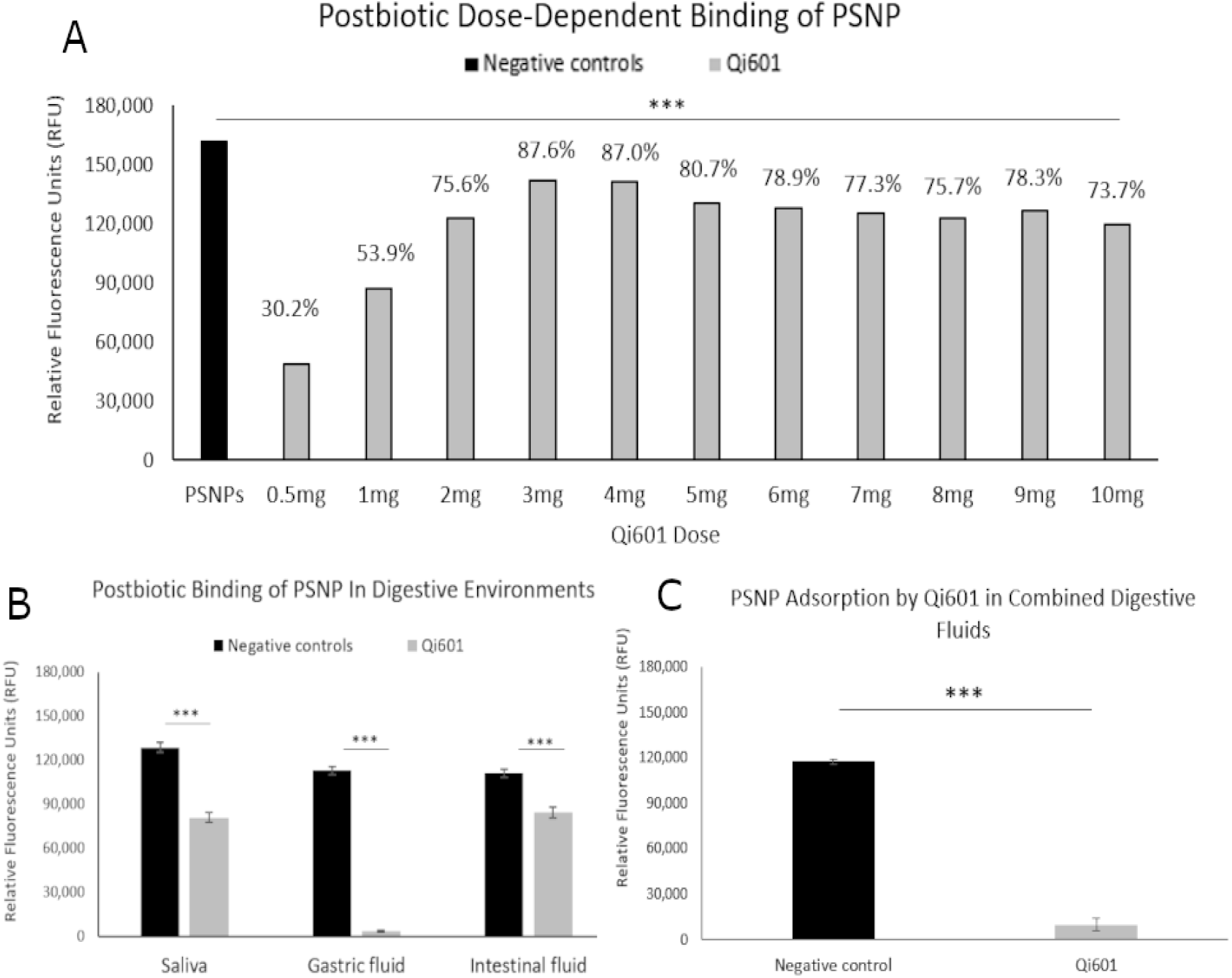
Qi601 rapidly binds PSNP in PBS and simulated digestive fluids. A) Baseline dose-response of Qi601-mediated PSNP adsorption in PBS, measured by residual PSNP fluorescence in the supernatant after separation of Qi601-associated material (n = 3 per group). B) Residual unbound PSNP fluorescence after incubation with Qi601 (3 mg/mL) in individual simulated digestive fluids, including salivary, gastric, and intestinal fluids (n = 9 per group). C) Residual unbound PSNP fluorescence after incubation with Qi601 (3 mg/mL) in a sequential simulated digestive-fluid model of gastrointestinal transit (n = 12 per group). Data are shown as mean ± SD. RFU, relative fluorescence units. *p < 0.05, **p < 0.01, ***p < 0.001, ****p < 0.0001.

Qi601 rapidly reduced unbound PSNP fluorescence in simulated digestive fluids. Significant reduction was detectable as early as 2 minutes in simulated salivary fluid. Adsorption varied by digestive condition, with lower adsorption under neutral-pH intestinal conditions and higher adsorption in simulated gastric fluid and in the sequential digestive-fluid model. Qi601 adsorbed PSNP in all tested conditions, with approximate adsorption of 37% in salivary fluid, 96% in gastric fluid, 24% in intestinal fluid, and 92% in the sequential saliva–gastric–intestinal model (all p < 0.0001; Fig. 1B and C).

To determine whether simulated digestion altered the ability of Qi601 to protect intestinal epithelial cells from PSNP exposure, Qi601 was subjected to sequential digestive-fluid exposure and then applied to Caco-2 monolayers maintained in digestive fluid diluted 1:2 in complete culture medium. Three exposure paradigms were tested: rescue, in which Qi601 was added after 24 hours of PSNP exposure; protection, in which Qi601 was added before PSNP exposure; and retention, in which pre-formed Qi601–PSNP agglomerates (QPAs) generated in digestive fluids were added to monolayers to assess whether bound PSNP remained retained within QPAs or were transferred to epithelial cells.

Rescue testing showed that Qi601 significantly reduced cell-associated PSNP even when applied after 24 hours of prior PSNP exposure (Fig. 2). This effect was time dependent. After 5 hours of Qi601 treatment, no significant reduction was observed in either surface-associated or intracellular PSNP signal (Fig. 2A and B). In contrast, 24 hours of Qi601 treatment significantly reduced surface-associated PSNP by 77% (*p* = 0.0002) and intracellular PSNP by 43% (*p* = 0.0014; Fig. 2C and D). Thus, Qi601 was not limited to a pre-exposure protective effect; it also reduced established epithelial PSNP burden after prior particle exposure, although this effect required longer treatment duration.

**Fig. 2.**
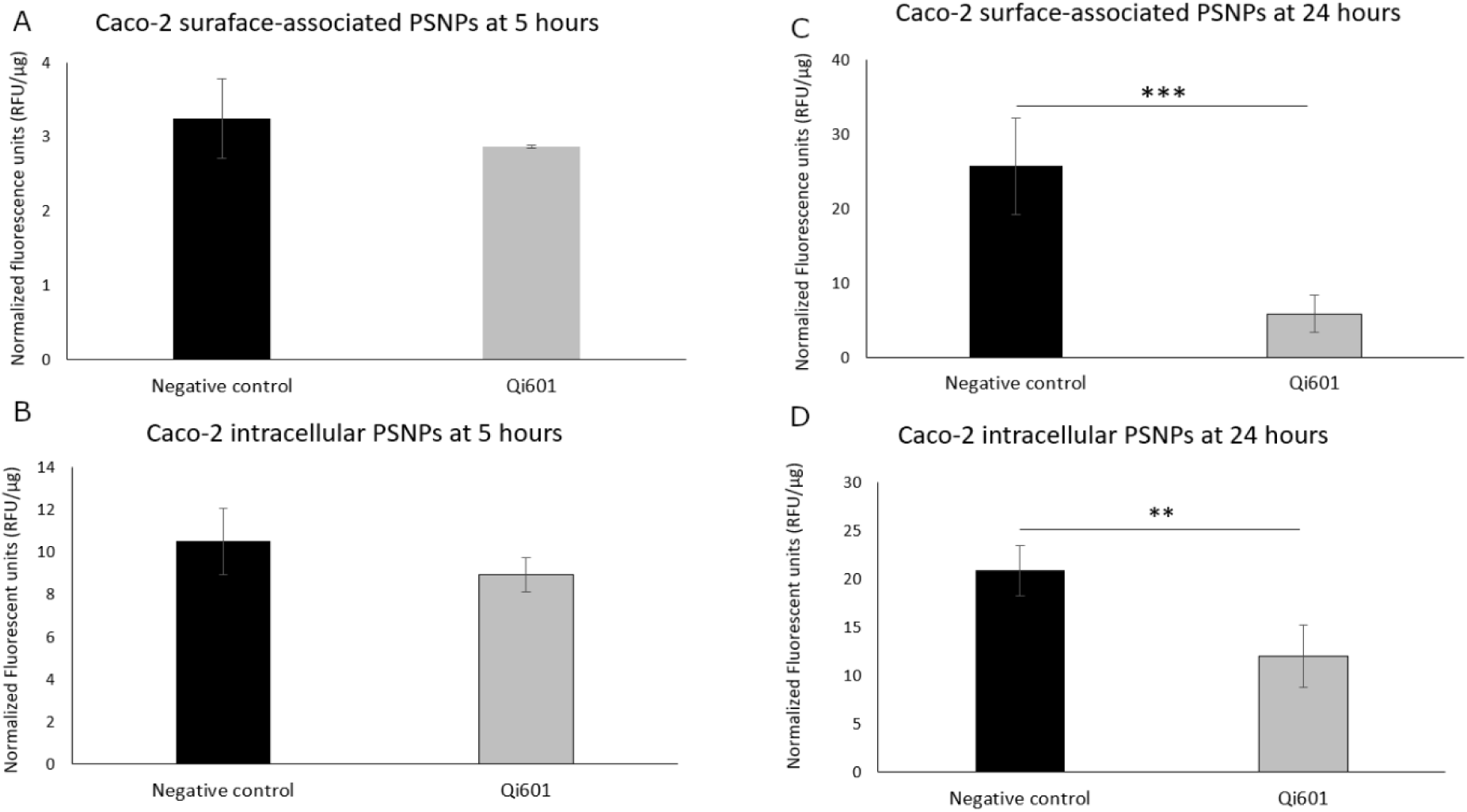
Qi601 reduces surface-associated and intracellular PSNP in Caco-2 cells after prior PSNP exposure. Caco-2 monolayers were pre-exposed to PSNP for 24 h, followed by treatment with Qi601 (3 mg/mL) or control medium for 5 h (A and B) or 24 h (C, D) (n = 5 per group). Surface-associated PSNP fluorescence (A and C) and intracellular PSNP fluorescence (B and D) were quantified and expressed as relative fluorescence units normalized to protein content per well (RFU/µg protein). Data are shown as mean ± SD. RFU, relative fluorescence units. * *p* < 0.05, ** *p* < 0.01, *** *p* < 0.001.

Prior studies in Caco-2 cells have shown that PSNP internalization is established at approximately 24 hours^27^, localizing predominantly to perinuclear areas and cytoplasmic vesicular compartments such as including vacuoles and lysosomes^28^. Therefore, after observing significant postbiotic-mediated reduction in intracellular PSNP burden, we used confocal laser scanning microscopy (CLSM) to evaluate the subcellular localization of PSNP internalized in pre-exposed Caco-2 cells.

Consistent with the preceding Caco-2 rescue assay, pixel-based quantitative confocal analysis demonstrated a Qi601-mediated reduction in intracellular PSNP signal in pre-exposed Caco-2 cells. After 24 hours of Qi601 treatment, significant reductions were observed in whole-cell PSNP fluorescence (p = 0.0015; Fig. 3A) and perinuclear PSNP fluorescence (p = 0.0018; Fig. 3C) compared with control, whereas cytoplasmic PSNP-associated fluorescence showed a smaller, non-significant decrease (Fig. 3B). These effects were not significant after 5 hours of Qi601 treatment in any measured compartment (Fig. 3A–C). Qi601 did not significantly reduce cytoplasmic PSNP signal at either time point, although this analysis may have been limited by high well-to-well variability and heterogeneous Caco-2 morphology in culture (Fig. 3B).

**Fig. 3.**
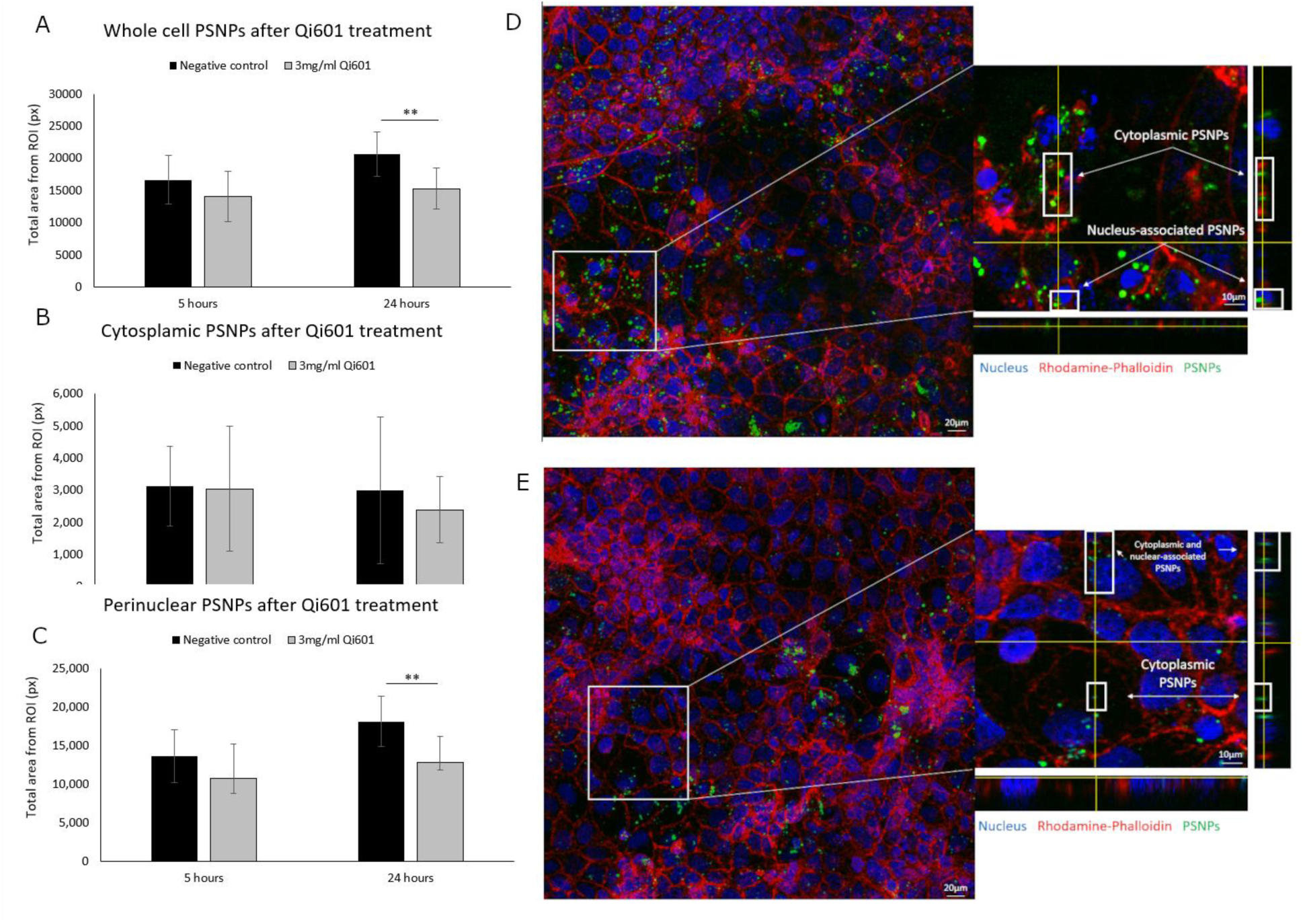
Qi601 reduces whole-cell and perinuclear PSNP fluorescence in a time-dependent manner. A–C) Whole-cell (A), cytoplasmic (B), and perinuclear (C) PSNP-associated fluorescence at 5 h and 24 h after Qi601 treatment. A total of 14–20 regions of interest (ROIs) were analyzed per condition. D) Representative confocal fluorescence micrograph of a Caco-2 monolayer exposed to PSNP for 24 h without Qi601 treatment. The left panel shows the full image used for analysis, and the right panel shows an orthogonal view from a Z-stack acquisition. Abundant PSNP fluorescence (green) is visible near the actin cytoskeleton/plasma membrane marker (rhodamine-phalloidin, red) and in perinuclear regions relative to nuclei stained with DAPI (blue). E) Representative confocal fluorescence micrograph of Caco-2 cells exposed to PSNP for 24 h followed by 24 h of Qi601 treatment. The left and right panels show reduced PSNP fluorescence, particularly in whole-cell and perinuclear compartments, compared with untreated control cells. PSNP-associated fluorescence was quantified and expressed as relative fluorescence units normalized to protein content per well (RFU/µg). Data are shown as mean ± SD. RFU: Relative Fluorescent Units. \**p* < 0.05, \*\**p* < 0.01, \*\*\**p* < 0.001

Representative CLSM images supported the quantitative findings. At 24 hours, untreated control cells showed abundant PSNP fluorescence throughout the cell, with prominent perinuclear accumulation consistent with intracellular redistribution after uptake (Fig. 3D). Qi601-treated cells showed an overall reduction in PSNP fluorescence, with markedly decreased perinuclear signal (Fig. 3E). Taken together, the rescue assay and CLSM analysis indicate that Qi601 reduces cell-associated and perinuclear PSNP burden even after epithelial PSNP exposure has already occurred.

Cellular protection assays were consistent with rapid-onset Qi601-mediated inhibition of initial PSNP association with Caco-2 monolayers and reduced subsequent intracellular PSNP accumulation. At 5 h, Qi601-protected cells showed 65% lower surface-associated PSNP fluorescence (p < 0.0001; Fig. 4A) and 67% lower intracellular PSNP fluorescence (p < 0.0001; Fig. 4C) compared with control cells. The protective effect diminished but remained significant at 24 h, when Qi601-treated Caco-2 cells showed 41% lower surface-associated PSNP fluorescence (p = 0.004; Fig. 4B) and 33% lower intracellular PSNP fluorescence (p = 0.021; Fig. 4D) compared with controls. Compared with the rescue assay, the protection assay showed earlier reductions in both surface-associated and intracellular PSNP fluorescence, whereas significant rescue effects were observed only after 24 h of Qi601 treatment.

**Fig 4.**
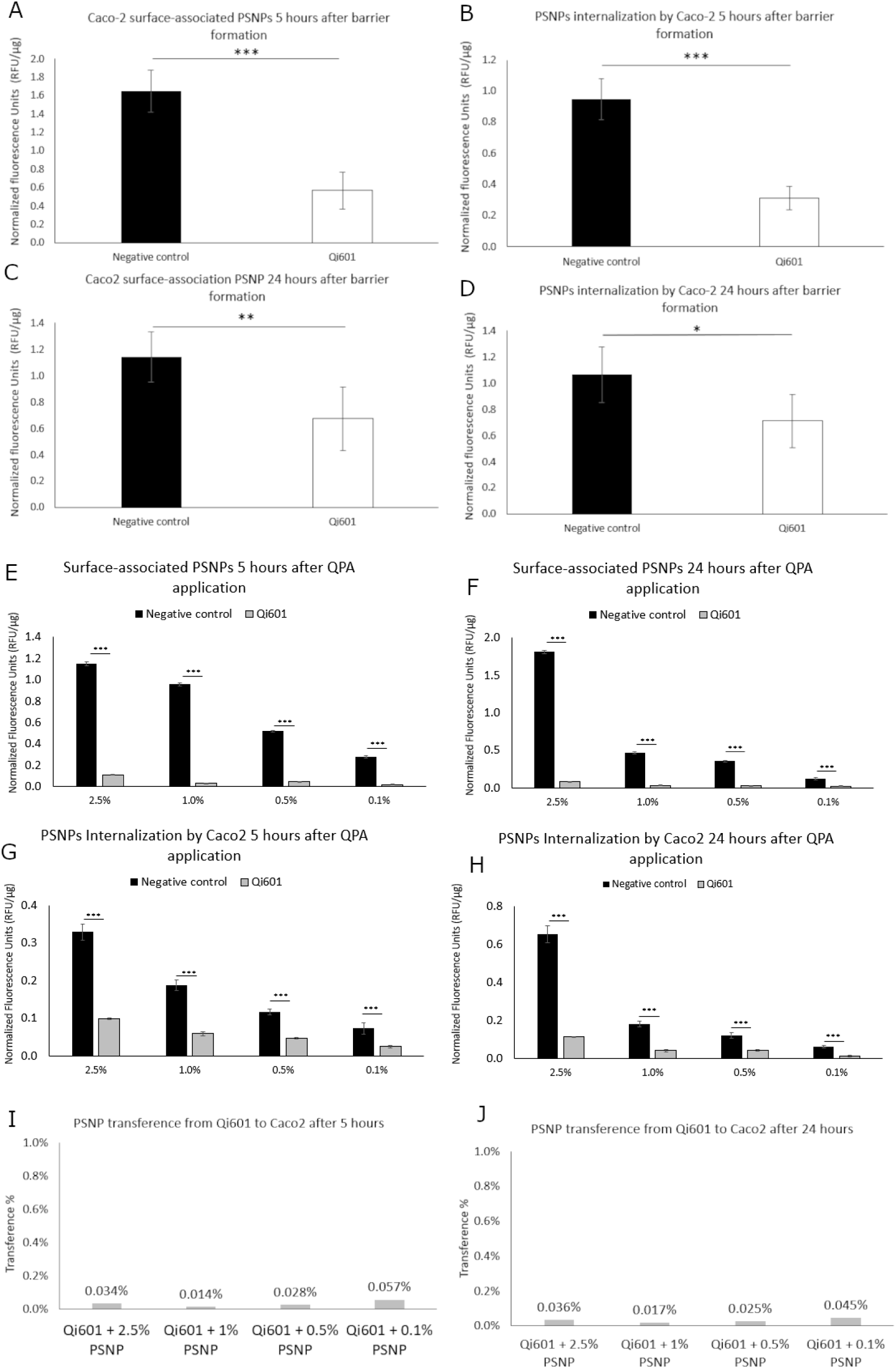
Qi601 forms a protective barrier against PSNP on Caco-2 cells with minimal transfer of postbiotic-bound PSNP to cells. A-D) Protection assay: Caco-2 monolayers were treated with Qi601 (3 mg/mL) prior to exposure to PSNP. Membrane-associated PSNP fluorescence (A, B) and cellular uptake (C, D) were quantified at 5 hours and 24 hours. E-H) Qi601–PSNP agglomerates (QPA) model: PSNP were pre-incubated with Qi601 to form QPAs prior to application to Caco-2 monolayers. Membrane-associated (E, F) and intracellular (G, H) PSNP fluorescence were quantified at 5 h and 24 h across a range of PSNP concentrations (0.1–2.5%). I-J) Transference (agglomerate) model: Percentage of PSNP fluorescence transferred from Qi601-associated complexes to epithelial cells at 5h and 24h. PSNP-associated fluorescence was quantified and expressed as relative fluorescence units (RFU/µg) normalized to protein content per well. Data are shown as mean ± SD. RFU: Relative Fluorescent Units. \**p* < 0.05, \*\**p* < 0.01, \*\*\**p* < 0.001

A critical requirement of the Qi601 sequestration model is that Qi601 retain bound PSNP rather than act as a carrier that delivers particles to the epithelium. Therefore, we next performed a transfer assay to determine whether PSNP pre-associated with Qi601 under sequential digestion conditions could redistribute from Qi601–PSNP agglomerates to adjacent Caco-2 monolayers. Compared with PSNP-only controls, Qi601–PSNP agglomerates markedly reduced both surface-associated and intracellular PSNP fluorescence across all tested concentrations and time points (*p* < 0.0001; Fig. 4E–H), consistent with stable sequestration of PSNP within Qi601-associated complexes. Complex retention appeared to be maintained over time, as transfer percentages were very low when calculated as total cell-associated fluorescence divided by total fluorescence associated with Qi601-adsorbed PSNP. Transfer ranged from 0.014% to 0.057% at 5 h and from 0.017% to 0.045% at 24 h, indicating minimal transfer of bound PSNP from Qi601 to epithelial cells (Fig. 4I and J).

The limited epithelial transfer of Qi601-bound PSNP suggested that PSNP remained associated with the postbiotic matrix rather than released to adjacent epithelial surfaces. To visualize Qi601–PSNP interactions directly, complementary imaging modalities were used. Fluorescence imaging showed dense PSNP signal colocalizing with discrete multicellular Qi601 clusters (Fig. 5A). AFM demonstrated PSNP arrayed along postbiotic surfaces with aggregated and organized topography not observed in controls (Fig. 5B). Confocal microscopy showed Qi601 clusters positioned above the epithelial monolayer, with PSNP signal predominantly localized to Qi601-associated material (Fig. 5C). Orthogonal Z-stack views showed reduced PSNP signal within the epithelial monolayer, supporting spatial separation between Qi601-associated PSNP aggregates and the epithelial cell layer. SEM demonstrated postbiotic agglomerates with densely adherent PSNP associated with Qi601 surface mesh-like material (Fig. 5D and E). Together, these imaging findings show close physical association between PSNP and Qi601 clusters, with limited localization of PSNP signal to the underlying epithelial monolayer.

**Figure 5.**
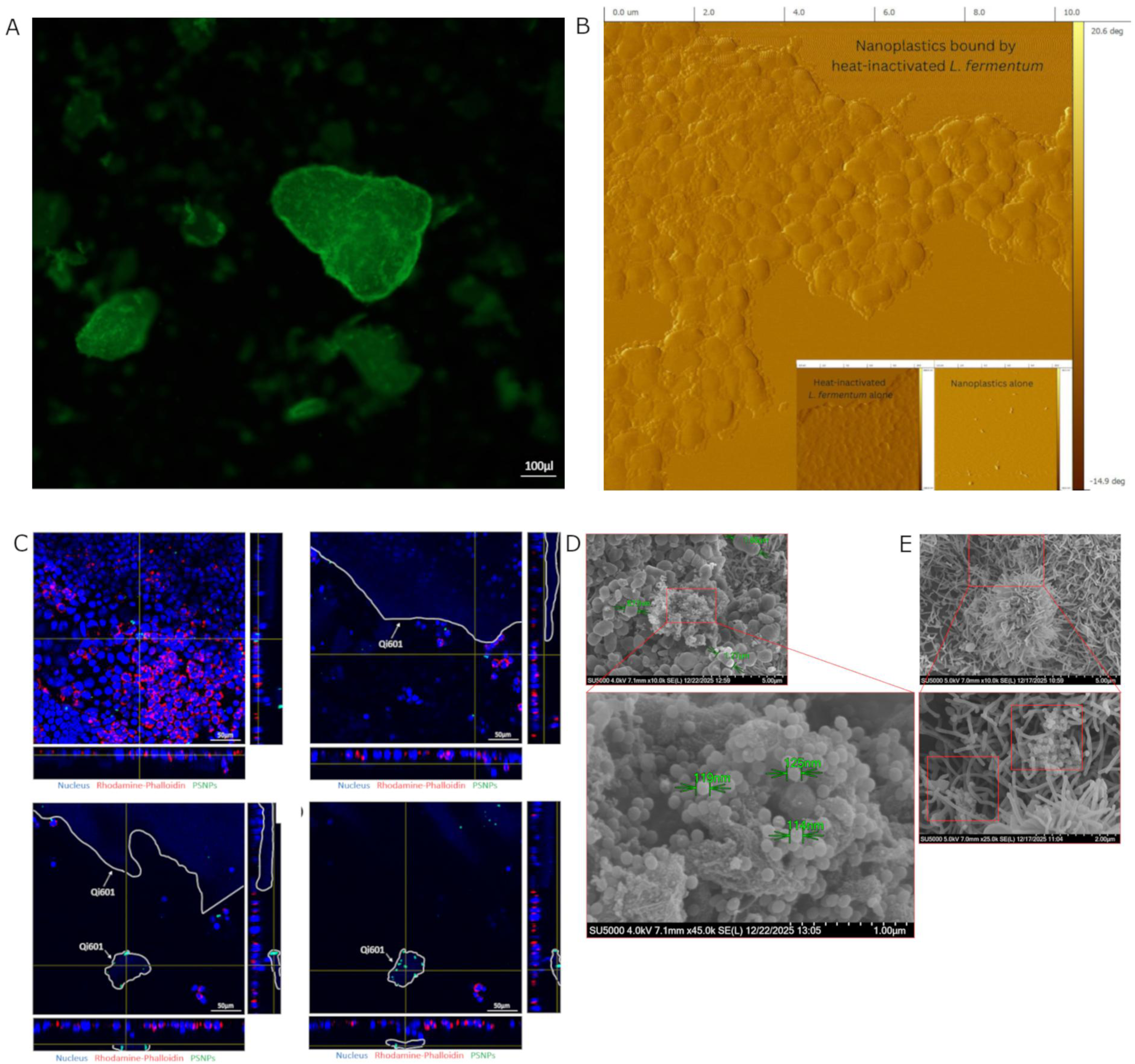
Multimodal imaging demonstrates Qi601–nanoplastic association, extracellular sequestration, and aggregate formation. (A) Fluorescence microscopy demonstrated dense PSNP signal associated with discrete Qi601 postbiotic assemblies, consistent with PSNP capture by Qi601 complexes. B) Atomic force microscopy (AFM) phase imaging showed PSNP organized along Qi601 surfaces, with an aggregated surface topography not observed in the PSNP-only control shown in the inset panel. C) Confocal laser scanning microscopy (CLSM) of Caco-2 monolayers exposed to PSNP in the presence of Qi601 showed Qi601 aggregates positioned above the epithelial monolayer. Orthogonal Z-stack views demonstrated separation between the base of a Qi601 aggregate (white circle) and the underlying Caco-2 cells. Multiple green-fluorescent PSNP foci were localized within smaller Qi601 aggregates, supporting extracellular sequestration of PSNP by Qi601. D) SEM imaging of Qi601-treated Caco-2 monolayers showed Qi601 aggregates on the epithelial surface with numerous PSNP clustered on or within the postbiotic material. Higher-magnification SEM demonstrated a mesh-like Qi601-associated matrix containing multiple PSNP and showed close association between PSNP and postbiotic surfaces. E) In the absence of Qi601, PSNP adhered directly to Caco-2 cells, with higher magnification showing numerous PSNP associated with epithelial microvilli. Scale bars are shown in each panel.

Having shown that Qi601 binds PSNP and limits epithelial PSNP exposure, we next examined whether this sequestration behavior could extend to a more heterogeneous oral microplastic exposure model. Recent studies have shown that mastication of chewing gum can release irregular polymeric fragments into saliva before swallowing^29^. Accordingly, chewing gum was selected as a proof-of-concept exposure model to qualitatively assess whether Qi601 could associate with gum-derived microplastics released during chewing.

Saliva samples collected before and after chewing commercially available gum demonstrated gum-derived microplastic particles in both control and Qi601-containing conditions. Nile Red staining with UV excitation^30^ revealed fluorescent particles consistent with polymeric material under both conditions. These particles showed irregular morphology and heterogeneous size distribution, with observed fragments ranging from approximately 20–100 µm, and occasional larger fragments extending to several hundred microns in this qualitative imaging model.

Qi601 appeared as variably sized multicellular structures and smaller postbiotic fragments (Fig. 6A). In Qi601-containing samples, postbiotic structures were observed in close association with gum-derived microplastic particles and, in some cases, appeared to bridge or aggregate multiple particles (Fig. 6B). Composite fluorescence images showed colocalization between Qi601 material and plastic-associated signal (Fig. 6C and D). The Qi601–plastic interface appeared broad and continuous rather than limited to isolated points of contact, consistent with extensive surface interaction between Qi601-associated material and the plastic fragments. Together, these findings indicate that Qi601 can associate with heterogeneous polymeric fragments generated during chewing.

**Figure 6.**
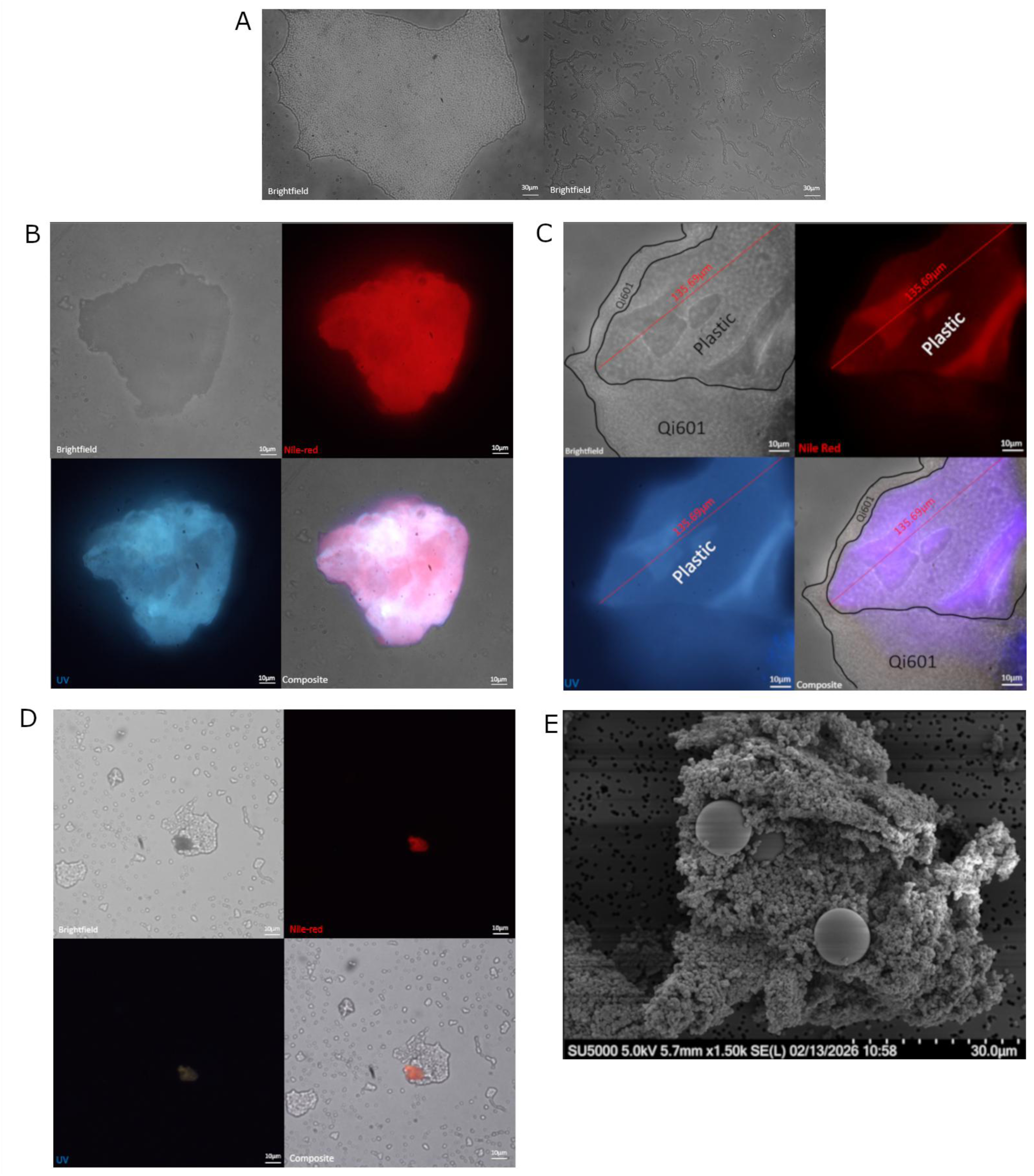
Microscopic visualization of chewing gum-derived microplastic particles extracted from saliva. A) Qi601 assemblies in PBS range from large aggregates of heat-killed biofilm made of hundreds of thousands of dead bacteria to smaller multi-cellular structures. B) Microplastic particles stained by Nile-red and fluorescent under UV light from the saliva of an individual who chewed commercial gum without Qi601. Background contains biological material such as epithelial cell debris, which do not fluoresce with Nile Red or UV light at the same intensity as synthetic polymer. Top left image: brightfield image; top right image: Nile red staining; bottom left image: fluorescent signals under UV light; bottom right image: composite. C) Microplastic particles are seen embedded in a large Qi601 postbiotic aggregate sampled from the saliva of an individual who chewed gum simultaneously with Qi601. Top left image: brightfield image; top right image: Nile red staining specifically stains the plastic particle; bottom left image: the plastic particle fluoresces blue under UV light; bottom right image: composite. The plastic particle is delineated within Qi601. D) Microplastic particles are seen embedded in a small Qi601 postbiotic aggregate sampled from the saliva of an individual who chewed gum simultaneously with Qi601. Top left image: brightfield image; top right image: Nile red staining specifically stains the plastic particle; bottom left image: the plastic particle fluoresces blue under UV light; bottom right image: composite. The plastic particle is delineated within Qi601 E) SEM image of a 10µm PSMPs embedded in a Qi601 aggregate.

**Fig. 7.**
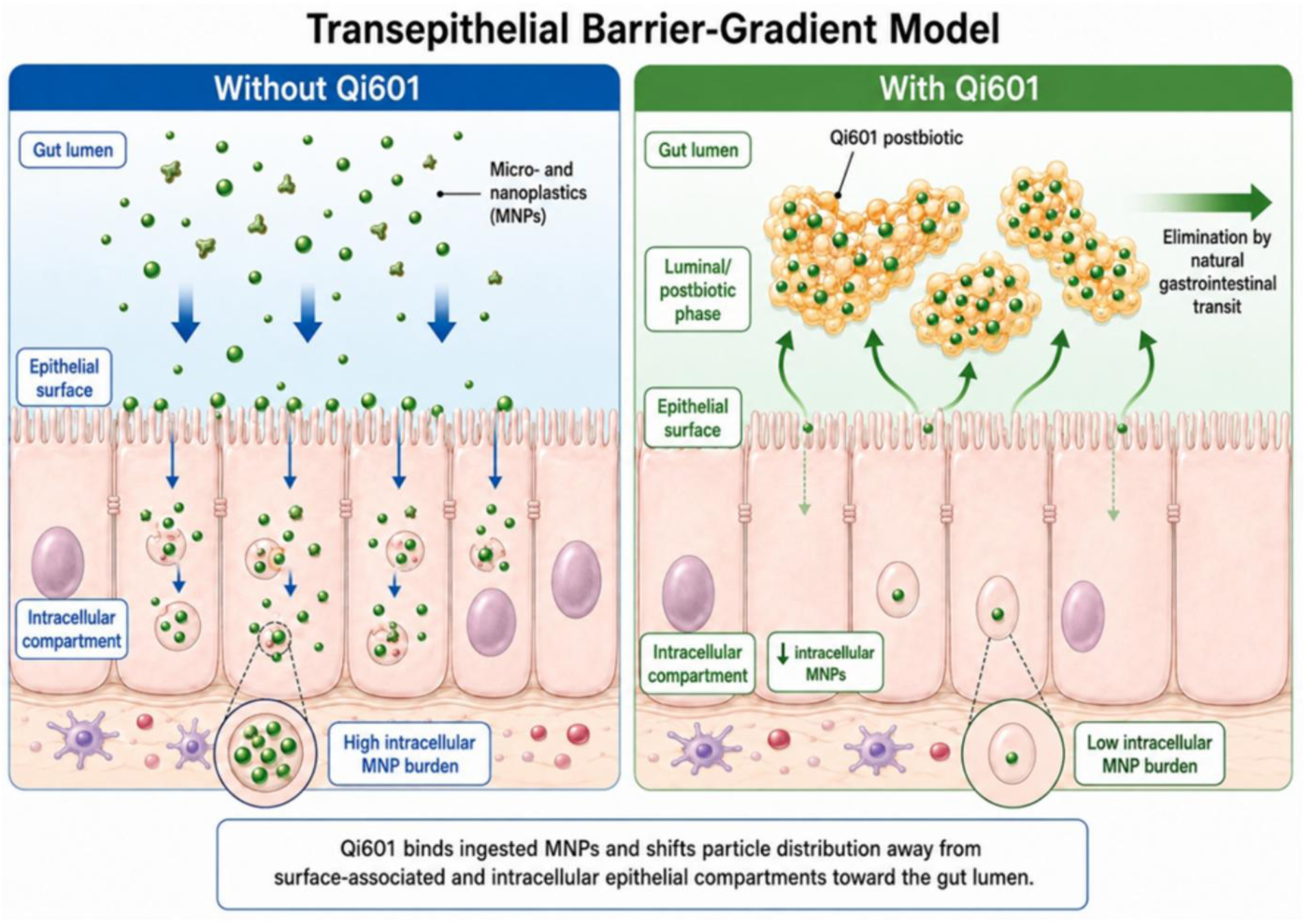
Qi601 transepithelial barrier-gradient model. Postbiotic Qi601 binds ingested MNPs to maintain them in the luminal compartment, where they can be available for elimination by natural gastrointestinal transit. The net effect is to reduce surface and intracellular MNP burden. Figure made in BioRender.com.

The gum-chewing samples extended the PSNP findings by demonstrating Qi601 association with larger, irregular plastic fragments generated under a human oral exposure condition. To determine whether Qi601 could also interact with microscale plastics under defined conditions, uniform 10-µm PSMP spheres were used as a controlled comparator. SEM showed close association between Qi601 and PSMP spheres, including apparent partial envelopment of particles by Qi601-associated material after incubation (Fig. 6E). This controlled microscale experiment supports the gum observations and indicates that Qi601–plastic interactions are not limited to nanoscale PSNP but extend to larger microscale plastic particles potentially encountered through the diet.

## Discussion

Together, confocal microscopy, rescue, and protection assays suggest that Qi601 may act through complementary mechanisms: “protective barrier” sequestration that limits early epithelial PSNP contact, and as a post-exposure sink effect that reduces established cell-associated and intracellular PSNP burden. The distinct kinetics of these effects provide further support for this dual-mechanism model and likely reflect differences in PSNP accessibility at the time of postbiotic introduction. In the protection assay, Qi601 was present during the initial exposure period, when PSNP were expected to remain largely extracellular or surface-associated and could therefore be rapidly sequestered before epithelial adhesion and uptake. This is consistent with the strong reduction in PSNP burden observed by 5 hours. In contrast, in the rescue assay, Qi601 was added after 24 hours of PSNP exposure, a time point at which prior Caco-2 studies have shown PSNP internalization and localization to perinuclear and vesicular compartments, including vacuoles and lysosomes.^27,28^

Because internalized particles are less accessible to extracellular Qi601, reduction of established PSNP burden likely depends on slower processes, such as dissociation of surface-bound PSNP, vesicular recycling or exocytosis, reduced re-entry, and ongoing extracellular sequestration.^31–34^ This model is consistent with the rescue assay, in which significant reductions in surface-associated and intracellular PSNP fluorescence were observed after 24 hours of Qi601 treatment but not after 5 hours. Together, the rapid protection effect, delayed rescue effect, CLSM reduction in perinuclear PSNP signal, and limited epithelial transfer of Qi601-bound PSNP support a model in which Qi601 creates a luminal PSNP sequestration sink, reducing the free PSNP fraction available for epithelial contact and uptake. To describe this effect, we use the term “transepithelial barrier gradient” to refer to a luminal-to-epithelial distribution shift in which Qi601 reduces the free PSNP fraction available at the epithelial interface and shifts measurable particle burden away from surface-associated and intracellular epithelial compartments toward the postbiotic/luminal phase.

Although the human health effects of MNP exposure remain incompletely defined, there is increasing interest in strategies that reduce human exposure to these foreign particles.^35,36^ Most plastic-remediation approaches have focused on environmental removal or degradation; however, approaches designed to reduce gastrointestinal absorption of ingested MPs are now beginning to receive attention.^37,38^ Given their safety profile, broad commercialization, ease of use, and direct relevance to diet-derived MP exposure, probiotic organisms have recently been explored as potential MNP biosorbents. Current evidence remains limited, but emerging studies have demonstrated strain-specific MP adsorption and increased fecal excretion of MPs in probiotic-treated mice.^43–46^

These findings extend, rather than contradict, recent probiotic-MNP studies. Teng et al. demonstrated that selected live probiotic strains can adsorb microplastics and enhance their excretion in vivo, supporting the general concept that microbial surfaces can act as gastrointestinal plastic-particle biosorbents. Shi et al. further showed that LAB-mediated mitigation of MNP toxicity depends not only on binding capacity but also on restoration of the gut environment, including barrier-related and microbiome-associated effects. Zhao et al. demonstrated that live *L. plantarum* ZP-6 can reduce PS-NP tissue retention, increase fecal excretion, and ameliorate inflammatory, barrier, and metabolic injury in murine models. In contrast, the present study focuses on postbiotic material and isolates the physical binding and epithelial-interaction component of this broader biological paradigm.

The principal distinction of Qi601 is that its proposed activity does not depend on live microbial viability, colonization, metabolic activity, or host-specific engraftment. Instead, Qi601 appears to act as a biofilm-derived particulate binding scaffold that retains PSNP under simulated digestive conditions and limits epithelial transfer of bound particles. This distinction may be important for product reproducibility, storage stability, and safety in heterogeneous human populations. Using orthogonal methods, the work here using a non-viable *L. fermentum* biofilm cellular mass demonstrates successful in vitro postbiotic-mediated adsorption of micro- and nanoplastics, including heterogeneous gum-derived microplastic fragments generated in the human mouth. Our results contrast directly with other studies, which have shown loss of MP-binding effect for heat-inactivated vs live probiotic strains. Though the work described here was not designed to be a mechanistic investigation, the loss of MP binding seen with non-viable lactobacilli in other studies may be related to the production of Qi601 as a pre-formed biofilm. It is well known that biofilms readily adhere to and colonize the hydrophobic surfaces of MPs in aqueous environments, called the “plastisphere”.^39^

Postbiotics may provide a safer and more controllable binding platform for luminal sequestration and elimination through normal gastrointestinal transit than probiotics.^40^ By definition, postbiotics are preparations of inanimate microorganisms and/or their components that confer a health benefit, and their development emphasizes defined microbial identity, inactivation procedures, confirmation of nonviability, and safety for intended use. Because microbial adsorption of plastic particles appears to be mediated largely by physical surface interactions, the microbial surface of Qi601 appears to be an organizing and durable scaffold for plastic particle capture.^15^ A nonviable postbiotic could therefore function as a luminal biosorbent, physically sequestering MNPs in the gut lumen and favoring fecal diversion, while avoiding risks and limitations associated with live microbial administration, including viability loss, microbiome engraftment, antimicrobial-resistance transfer, unpredictable host colonization and plastic-mediated toxicity to the probiotic organisms themselves.^41^ Use of a postbiotic may also be particularly attractive in heterogeneous human populations, where product stability, reproducible dosing, and safety are central translational requirements.

Use of postbiotics may also reduce the theoretical risk of MP “shuttling” associated with live probiotics. Microplastics can adsorb and transport chemical co-contaminants, including metals and organic pollutants, and contaminant release may vary with local physicochemical conditions.^42^ Live probiotic organisms can also adhere to mucus and epithelial surfaces through strain-specific adhesion mechanisms.^43^ Together, these observations raise the theoretical possibility that a live MP-binding organism could persist at the mucus–epithelial interface long enough to redistribute MP-associated chemicals or particles. In contrast, our transfer-assay results indicate durable binding between Qi601 and PSNP, with minimal transfer of bound particles to gut epithelial surfaces. A nonviable postbiotic designed to preferentially bind MPs without colonizing or functioning as a potential vector for delivery to human cells may therefore be better suited as a transient gut sequestrant.

Consistent with the concept that live bacterial viability is not required to reduce epithelial nanoplastic internalization, heat-treated *L. delbrueckii* subsp. *bulgaricus* 2038 and *S. thermophilus* 1131 were recently shown to suppress PSNP internalization in differentiated Caco-2 monolayers.^44^ However, the authors found no evidence of direct LAB-PSNP aggregation and instead suggested that soluble bacterial cell components mediated an epithelial effect. To our knowledge, this is the first report demonstrating direct postbiotic–MNP binding under simulated human digestive conditions, supported by epithelial transfer-stability testing and human oral proof-of-concept imaging. To our knowledge, this is also the first microbial-derived MNP-binding study to include a post-exposure epithelial rescue model showing reduced established surface-associated and intracellular PSNP burden after prior epithelial exposure.

Prior probiotic studies have demonstrated that selected live LAB strains can adsorb micro- and nanoplastics and mitigate toxicity in vivo. The present study extends this paradigm to a biofilm-derived postbiotic and provides direct evidence of binding under simulated digestion, reduced epithelial interaction in protection and rescue models, minimal epithelial transfer of postbiotic-bound particles, multimodal visualization of postbiotic–plastic association, and first-in-human oral proof of concept. Together, these findings support a “transepithelial barrier-gradient” model in which Qi601 physically binds ingested nano- and microplastics, reducing surface-associated and intracellular epithelial burden while favoring retention within the gut lumen, where bound particles can undergo elimination through normal gastrointestinal transit.

### Limitations and Future Studies

This study has several limitations. First, although the study was designed to evaluate postbiotic Qi601 as a gastrointestinal MP sequestrant, the primary plastic used was polystyrene. Polystyrene remains one of the most commercially available and well-characterized model nanoplastics, and while useful for in vitro studies, PSNP do not capture the full polymeric, chemical, morphologic, and size heterogeneity of human-relevant micro- and nanoplastics. Furthermore, to generate detectable signal, the amount of nanoplastics used in these experiments far exceeds any dietary MNPs encountered in the human diet. Second, although Caco-2 monolayers are a standard intestinal epithelial model, they do not incorporate important in vivo gastrointestinal features, such as immune cells, peristalsis, microbiota, complex intestinal architecture, and real-world food and nutrient exposures.

We attempted to address these limitations by testing Qi601–PSNP binding in a physiologically relevant human digestion model, and in a proof-of-concept chewing gum study. To our knowledge, the chewing-gum experiment described here is the first human oral proof-of-concept study evaluating direct binding between a proposed MNP-binding intervention and gum-derived microplastic fragments under mastication conditions. However, it does not evaluate capture efficiency, dose-response behavior, or reduction in ingested plastic burden. Finally, unlike recent probiotic studies, this work did not measure fecal MNP excretion, biodistribution, molecular epithelial responses or microbiome remodeling. Therefore, the present findings should be interpreted as evidence of postbiotic binding, epithelial exposure reduction, and oral proof of concept rather than proof of reduced whole-body MNP burden.

Future studies are underway to quantify Qi601 binding across additional common polymers, incorporate more sophisticated biological systems such as gut organoids and animal models, define the physicochemical determinants of Qi601 binding, and evaluate inflammatory, oxidative-stress, washout, and re-dosing endpoints to determine whether replenishing Qi601 can sustain protection over longer exposures. Ultimately, these findings support the translational rationale for Qi601 as a nutritional supplement candidate for microplastic mitigation, while defining the next in vivo studies to evaluate Qi601-mediated gastrointestinal sequestration of ingested MNPs.

## Supplemental Material

Video 1: A time-lapse video showing how QPA of Qi601 and 0.2 μm red dye nanoplastics spheres settle at the bottom of a 15 ml conical tube within 4-5 minutes, while the negative control that lacks Qi601 shows the red dye nanoplastics remain in suspension.

